# Immunoglobulin G binding as a quantitative marker of hepatocellular death across acute and chronic liver injury

**DOI:** 10.1101/2025.11.27.690787

**Authors:** Lukas Neckermann, Pia Erdösi, Anne Dropmann, Amnah Othman, Johanna Weinert, Jan Albin, Stephanie D. Wolf, Brigitte begher-Tibbe, Iris von Recklinghausen, Tiziana Caccamo, Matthias P. Ebert, Johannes G. Bode, Jan Hengstler, Steven Dooley, Seddik Hammad

## Abstract

**Background & Aims:** Accurate detection of hepatocellular death is fundamental for understanding liver injury, intoxication, regeneration, and fibrosis. Conventional markers, such as serum transaminases and histopathological scoring, suffer from limited temporal resolution, high variability, and observer dependence. We evaluated immunoglobulin G (IgG) binding as a quantitative and spatially resolved marker of hepatocyte death in acute and chronic liver injury models.

**Methods:** Male C57BL/6 mice were subjected to acute carbon tetrachloride (CCl₄) intoxication (1600 mg/kg, single dose), dose-escalation (0-800 mg/kg), and chronic injury paradigms including Western diet (WD), WD+CCl₄, and Mdr2-/- mice with or without a single CCl₄ challenge. The serum ALT and AST levels were measured. Liver sections were stained with IgG, Hematoxylin and Eosin (H&E), bromodeoxyuridine (BrdU), glutamine synthetase (GS), CD26, and alpha-smooth muscle actin (Acta2). Spatial and integrative transcriptomic analyses were performed to characterize the IgG⁺ hepatocyte dead regions.

**Results:** Hepatocellular IgG labeling emerged as early as 6h post-CCl₄, peaked at 72–96h, and declined during regeneration. IgG-positive areas correlated strongly with Ishak necroinflammatory score (r=0.70) and serum transaminase levels (p=0.74-0.85), surpassing both in Receiver Operating Characteristic (ROC) analyses (AUC=0.92-0.95). IgG bound to both apoptotic (TUNEL⁺) and necrotic (TUNEL⁻) hepatocytes. In chronic liver injury models, IgG deposition was localized to the injury zones and correlated with ALT/AST, irrespective of etiology. Multiplex imaging revealed IgG-positive necrotic cores surrounded by proliferating hepatocytes and Acta2^+^ myofibroblasts. Spatial transcriptomics identified immune cell enrichment, FcγR-mediated signaling, phagocytosis, and vascular remodeling within the IgG-marked regions.

**Conclusions:** IgG immunostaining provides a robust, quantitative, and pathologist-independent readout of hepatocellular death, which scales with injury severity, delineates necrotic zones, and reveals immune-active microenvironments. These findings establish IgG-based detection as a versatile, high-resolution tool for assessing liver injury, regeneration, and fibrosis.

**Conflict of Interest declaration:** The authors declare that they have no affiliations with or involvement in any organization or entity with any financial interest in the subject matter or materials discussed in this manuscript.

**Graphical abstract:** 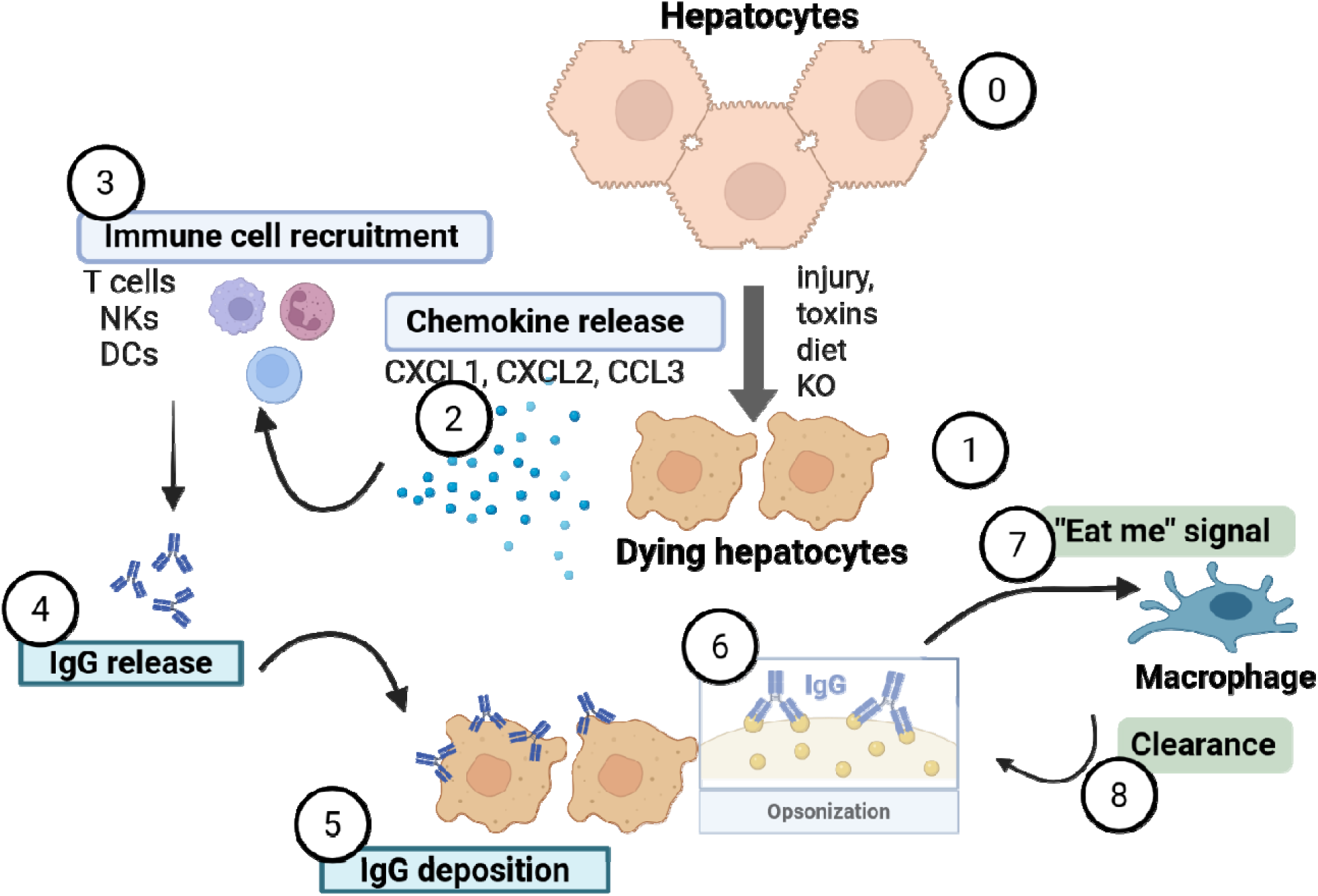

Liver injury induced by toxins, dietary stress, or genetic knockout provokes chemokine release (i.e. CXCL1, CXCL2, and CCL3) and recruitment of immune cells, including T cells, NK cells (NKs), and dendritic cells (DCs). Activated immune cells secrete IgG, which binds to damaged hepatocytes, leading to IgG deposition and opsonization. Opsonized hepatocytes expose “eat me” signals, promoting their phagocytic clearance and contributing to the resolution of liver injury and restoration of hepatic homeostasis.

## Introduction

The liver serves as a central hub for metabolic detoxification and immune surveillance with unique regeneration capability, rendering it particularly vulnerable to insults from xenobiotics, viral infection, metabolic overload, and cholestatic injury (1, 2). Hepatocellular death is a central and driving event in both acute and chronic liver diseases, including drug-induced liver injury (DILI), acute liver failure (ALF), metabolic dysfunction-associated steatohepatitis (MASH), cholestatic fibrosis, parenchymal injury initiation, inflammatory activation, and fibrogenic progression. This process is recognized as a key trigger for the amplification of inflammation and development of fibrosis across diverse etiologies of liver injury (3–7). The magnitude, mode (apoptosis, necrosis, necroptosis, and pyroptosis), and spatial context of hepatocyte death play a critical role in determining whether the liver successfully regenerates and returns to homeostasis or progresses to chronic injury, fibrogenesis, and organ failure. Different forms of hepatocyte death initiate distinct inflammatory signaling pathways and immune responses, which can either promote effective tissue repair or drive persistent injury and fibrosis, ultimately affecting the liver’s capacity for regeneration (6, 8). Thus, accurately detecting both the magnitude and spatial context of hepatocyte death is essential for predicting the disease trajectory and guiding targeted interventions to preserve liver function.

Despite the centrality of hepatocyte death in liver pathophysiology, current tools for its detection and quantification remain limited. Serum aminotransferases, specifically alanine aminotransferase (ALT) and aspartate aminotransferase (AST), are the most commonly used indirect biomarkers for assessing hepatocellular injury; however, their presence in the blood primarily reflects enzyme leakage from hepatocytes rather than direct cell death (9, 10). These enzymes have relatively long circulating half-lives, which diminish their temporal resolution and the ability to indicate immediate or ongoing injury (10, 11). ALT and AST blood levels are also affected by substantial inter-individual variations and can fluctuate widely within the same disease process, complicating their reliability, especially in chronic or slowly progressing liver disease settings (12, 13). Histopathology remains the gold standard for assessing tissue injury and necroinflammation with scoring systems such as Ishak or METAVIR in chronic liver disease (14, 15). However, these scores are inherently semi-quantitative, dependent on pathologist interpretation, and susceptible to sampling bias with limited temporal resolution, thereby constraining their applicability in biomarker development and mechanistic studies (16, 17). Accordingly, there is an urgent need for quantitative, reproducible, mechanistically and spatially resolved biomarkers that accurately capture hepatocellular death and its microenvironmental consequences.

One biologically plausible but under-exploited route is via the binding of circulating immunoglobulin G (IgG) to hepatocytes, whose plasma membranes become permeabilized during death processes. Loss of membrane integrity is a unifying terminal event in most hepatocellular death pathways, enabling the exposure of intracellular antigens, uptake of serum proteins, and engagement of immune clearance pathways. IgG is synthesized by plasma cells as a fundamental component of the adaptive immune response (18, 19), ass the most abundant immunoglobulin in human serum (∼75%) (20). It circulates ubiquitously throughout the intravascular and extravascular compartments, rendering it immediately available for immune surveillance (18, 21, 22). In this capacity, IgG functions as a potent opsonin for apoptotic and necrotic cells (23). The Fc region of the antibody specifically engages Fc gamma receptors (FcγRs) on phagocytes, thereby facilitating the recognition and clearance of cell debris via FcγR-mediated phagocytosis (26). Moreover, a recent mechanistic study demonstrated that natural IgG and IgM enhance the clearance of necrotic debris after acute acetaminophen liver injury, thereby facilitate regeneration (27). Studies on systemic autoimmune diseases have shown that IgG deposition within the organ parenchyma can promote inflammation and tissue damage (27, 28). Moreover, tissue imaging and spatial transcriptomics suggest that immunoglobulin deposition can reveal immune-active spots (find-me or eat-me signals) and necroinflammatory niches within the injured liver (27, 29, 30). Despite this, the potential of IgG binding as an intrinsic and quantifiable indicator of hepatocyte death has not yet been systematically explored in acute and chronic models of liver injury.

Here, we investigated IgG binding as a quantitative and spatially resolved marker of hepatocellular death in acute and chronic liver injury paradigms. Using CCl₄-induced acute toxicity in time and dose escalation models, as well as chronic injury settings including Western diet (WD) feeding, Mdr2-/- mice, and combined CCl₄ exposure, and we systematically compared IgG labeling to conventional markers such as serum transaminases and histopathological Ishak scoring. Furthermore, we integrated multiplex imaging and spatial transcriptomics to delineate the immune, vascular, and regenerative microenvironments surrounding the IgG-positive necrotic zones. Our findings established IgG immunostaining as a quantitative, reproducible, and pathophysiologically informative biomarker of hepatocellular death that bridges histological and molecular readouts across diverse models of liver injury.

## Material and Methods

### Animals and experimental design

Male C57BL/6 mice aged 8-12 weeks and Mdr2-deficient mice (Mdr2-/-) with a BALB/c genetic background (60 weeks old) were used in this study. All mice were housed under specific pathogen-free (SPF) conditions in a controlled environment with a 12-hour light-dark cycle and had ad libitum access to standard chow and water. All procedures followed the national and international guidelines for animal welfare, and were conducted in accordance with the Declaration of Helsinki.

### CCl₄-induced liver injury in C57BL/6 mice

In the time-course study, C57BL/6 mice (n=4–5 per time point) received a single intraperitoneal injection of CCl₄ (1600 mg/kg) and were sacrificed 1, 3, 6, 12, 24, 48, 72, 96, 144, and 384-hours post-injection to evaluate the temporal progression of liver injury, including acute hepatocellular damage, inflammation, and subsequent regenerative or fibrotic changes. In a dose-dependent study, male C57BL/6 mice (n=4 per dose) received a single intraperitoneal injection of CCl₄ at 25, 50, 100, 200, 400, or 800 mg/kg, and were sacrificed 48h later to assess IgG levels in relation to dose-dependent liver injury. The study protocol was approved by the institutional ethics committee (#35-9185.81/G-15/24 and #35-9185.81/G-223/20).

### CCl₄-induced liver injury in aged Mdr2-/- mice

Aged Mdr2-/- mice (n = 6 per time point, three males and three females) received a single intraperitoneal injection of CCl₄ (2400 mg/kg) and were sacrificed 24, 48, and 96h post-injection. Age-matched untreated Mdr2-/- (Mdr2KO) mice (n=8; four males, four females) were sacrificed without treatment to assess the baseline phenotype. The study protocol was approved by the institutional ethics committee (#35-9185.81/G-24/20 and #35-9185.81/G-22/20).

### Western diet (WD) model

Male C57BL/6J mice (n=3) were maintained on standard chow diet for 8 weeks. At 8 weeks of age, mice were switched to a Western diet (composition: 20% fat, 45% carbohydrate, 22% protein) supplemented with 23.1 g/L fructose and 18.9 g/L glucose in drinking water and injection of corn oil for 15 weeks. Mice were sacrificed at 24 weeks of age to assess diet-induced metabolic dysfunction-associated steatotic liver disease (MASLD).

### Western diet plus CCl₄ (WD+CCl₄) model

Male C57BL/6J mice (n = 6) were fed WD and additionally 23.1 g/L fructose and 18.9 g/L glucose in their drinking water from the age of 8 weeks. In addition, they received weekly intraperitoneal injections of CCl₄ (0.32 µg/g body weight). Mice were sacrificed at 24 weeks of age (after 15 weeks of combined Western diet and CCl₄ treatment) to assess the synergistic effects of metabolic stress and chronic hepatotoxic injury on liver pathology, fibrosis development, and tumorigenesis.

### Tissue collection and processing

At designated time points, mice were anesthetized, and blood was collected prior to organ harvest. The left lobe of the liver was fixed in 4% paraformaldehyde (#P087.1, Karlsruhe, Germany) in phosphate Buffered Saline (PBS) for histopathological analysis.

### Clinical Chemistry

Blood was collected via the retro-orbital venous plexus and centrifuged at 14,000 g for 3 min at 4°C. Plasma was transferred to a fresh tube and stored at –20°C until analysis. Plasma levels of alanine aminotransferase (ALT) and aspartate aminotransferase (AST) were measured using an automated Hitachi analyzer.

### Hematoxylin and Eosin (H&E) staining

To score necrotic regions using Ishak scoring, formalin-fixed, paraffin-embedded (FFPE) liver tissue sections were stained with hematoxylin and eosin (HE) according to standard protocols. The sections were deparaffinized by three successive washes in Roti-Histol (10 min each) and rehydrated using a graded isopropanol series (100%, 96%, 90%, and 70%; 5 min per step). After rehydration, sections were rinsed in distilled water for 5 min and incubated with Mayer’s hematoxylin for 5 min. Following nuclear staining, the slides were rinsed under running tap water for 15 min. Counterstaining was performed using 4% eosin for 4 min, followed by a brief rinse in distilled water. Dehydration was carried out using a graded isopropanol series, including two final washes in 100% isopropanol (5 min each). The slides were mounted and scanned using an Aperio 8 Slide Scanner (Leica Microsystems, Wetzlar, Germany).

### Histological evaluation

Histological evaluation of the liver samples was performed using the Ishak scoring system (15), which assesses both necroinflammatory activity and fibrosis stage. Necroinflammatory activity was scored on a scale from 0 to 18 based on lobular inflammation. For each section, ten randomly selected fields of view were assessed at 200× magnification (20× objective).

### IgG-staining protocol

FFPE liver tissue sections (4µm) were deparaffinized in xylene and rehydrated through a descending ethanol series, as described for the hematoxylin and eosin (H&E) staining protocol, with modifications (31). Heat-induced epitope retrieval (HIER) was carried out in Tris-HCl buffer (1.92 g/L citric acid, 500 µL/L Tween-20, pH 6.0) using a microwave with cycled heating (15s on, 45s off) for 10 min following the initial boiling phase. To minimize nonspecific antibody binding, the sections were incubated with 3% bovine serum albumin (BSA) in phosphate-buffered saline (PBS) for 1h at room temperature. Subsequently, the slides were incubated with Cy3-conjugated donkey anti-mouse IgG secondary antibody (#715-166-150; Jackson ImmunoResearch, PA, USA) diluted 1:100 in 0.3% BSA for 1 h at room temperature. After washing, the sections were mounted using Fluoroshield mounting medium containing DAPI (#F6182; Sigma-Aldrich, MO, USA) to visualize the nuclei. Fluorescent images were acquired using a confocal laser-scanning microscope (SP8; Leica Microsystems, Wetzlar, Germany) or FV1000 (Olympus, Tokyo, Japan).

### IgG positivity quantification

IgG positivity in the liver tissue was quantified using immunofluorescence confocal images and Fiji (ImageJ) software. Multichannel IF images (2000 × 2000 × px) were first split into individual channels, and the IgG-specific channel (e.g., Alexa Fluor 547 or Cy3) was selected for analysis. The IgG signal was filtered to suppress unspecific signals (median filter) and thresholded using an automated method (max. Entropie) to generate a binary mask, representing the IgG-positive areas. For normalization, the tissue area was defined using the DAPI channel as the nuclear marker. The percentage of IgG-positive areas was calculated relative to the total tissue area, and the mean fluorescence intensity was recorded for each image.

### Co-immunofluorescence staining

FFPE liver sections were deparaffinized in xylene and rehydrated using a graded ethanol series. Heat-induced epitope retrieval (HIER) was performed using either EDTA or Tris-NaCitrate-Dihydrate buffer, selected according to the specific antibody requirements (31). The primary antibodies were applied individually or in combination for immunofluorescence staining. The antibodies and their respective dilutions were rabbit anti-α-SMA (Acta2; #ab5698, Abcam, 1:100), rat anti-BrdU (#MCA2060, AbD Serotec, 1:500), goat anti-mouse DPPIV/CD26 (#AF954, R&D Systems, 1:100), and rabbit anti-glutamine synthetase (#G2781, Sigma, 1:2000). The primary antibodies were diluted in 0.3% BSA in PBS and incubated overnight at 4°C. After thorough washing, the sections were incubated for 1 h at room temperature with the corresponding fluorescently labeled secondary antibodies: Cy3-conjugated donkey anti-mouse IgG (#715-166-150, Jackson ImmunoResearch, 1:200), Alexa Fluor 647-conjugated donkey anti-mouse IgG (#715-606-150, Jackson ImmunoResearch, 1:500), Cy3-conjugated donkey anti-rabbit IgG (#711-166-152, Jackson ImmunoResearch, 1:200), Alexa Fluor 488-conjugated donkey anti-rat IgG (#712-546-150, Jackson ImmunoResearch, 1:100), and Alexa Fluor 488-conjugated donkey anti-goat IgG (#705-546-147, Jackson ImmunoResearch, 1:100). The nuclei were counterstained with DRAQ5 (#4084, Cell Signaling, 1:1000) for 5 min. Sections were mounted using Fluoroshield mounting medium containing DAPI (#F6182, Sigma-Aldrich) to preserve fluorescence and allow for nuclear visualization. Imaging was performed using confocal microscopy with Leica TCS SP8, Aperio Slide Scan 8 (Leica Microsystems, Wetzlar, Germany), and Olympus FV1000 (Olympus Corporation, Tokyo, Japan), providing high-resolution images for cellular and subcellular structure analysis.

### Dead cell quantification by TUNEL and IgG staining

Apoptotic cells were quantified using the In Situ Cell Death Detection Kit, Fluorescein (#11684795910, Roche) according to the manufacturer’s instructions. Antigen retrieval was performed in Tris-NaCitrate-Dihydrate buffer at 95°C for 60 min, followed by a 10-minute incubation with ready-to-use Proteinase K (#S3020, Dako) to permeabilize the tissue. The TUNEL reaction mixture was prepared according to the manufacturer instructions. For colocalization studies, Cy3-conjugated donkey anti-mouse IgG (#715-166-150, Jackson ImmunoResearch, 1:100) was added to the TUNEL reaction mixture. The sections were incubated for 60 min at 37°C in a humidified chamber followed by thorough washing. Sections were mounted using Fluoroshield mounting medium containing DAPI (#F6182, Sigma-Aldrich) to preserve fluorescence and allow for nuclear visualization. Imaging was performed using a Leica TCS SP8 confocal microscope, enabling high-resolution detection of apoptotic cells and colocalized antibody signals.

### Macrophage visualization by F4/80 immunohistochemistry

To visualize macrophages, F4/80 antibody staining was performed as previously described (4, 32) with slight modifications. Paraffin-embedded tissue sections were deparaffinized in xylene and rehydrated in a graded ethanol series. Antigen retrieval was performed by incubating the sections with ready-to-use Proteinase K (#S3020, Dako) for 10 min. Endogenous peroxidase activity was blocked using the Dako REAL Peroxidase-Blocking Solution (#S202386-2, Dako, ready-to-use). The sections were incubated overnight at 4°C with rat anti-F4/80 antibody (#MCA497, Bio-Rad, 1:100), followed by a 30-min incubation at room temperature with biotinylated anti-rat IgG (#BA-9400, Vector Labs, 1:2000). Subsequently, sections were incubated with streptavidin-conjugated horseradish peroxidase (#016-030-084, Jackson ImmunoResearch, 1:250) for 30 min at room temperature. Staining was visualized using diaminobenzidine (DAB), and the sections were counterstained with hematoxylin. The slides were mounted and scanned using an Aperio 8 Slide Scanner (Leica Microsystems, Wetzlar, Germany) to obtain high-resolution digital images.

### Affymetrix gene array analysis

Publicly available transcriptomic data from mouse liver samples following CCl_4_-induced injury were retrieved from ArrayExpress (E-MTAB-2445, https://www.ebi.ac.uk/biostudies/arrayexpress (33). The dataset includes a time-course of liver samples collected at defined intervals post-CCl₄ administration. Raw data were processed and normalized according to standard protocols, and downstream analyses were performed to assess gene expression dynamics across the injury and recovery phases.

### GeoMx digital spatial profiling (DSP) for spatial transcriptomics

Spatial transcriptomic analysis was performed using GeoMx Digital Spatial Profiler (NanoString) with probes from the Mouse Whole Transcriptome Atlas (Version 1). FFPE tissue sections (5μm thickness) were baked at 60°C for 30 min, followed by deparaffinization in xylene. Antigen retrieval was performed in Tris-EDTA buffer (pH 9.0, #00-4956-58; Thermo Fisher Scientific) for 20 min, and sections were subsequently treated with 1 μg/mL Proteinase K (#AM2546; Thermo Fisher Scientific) for 15 min at 37°C. A 5-min fixation step with 10% neutral buffered formalin (#15740-04; EMS Diasum) was performed prior to RNA probe hybridization, which was carried out overnight at 37°C. After stringent washing, the sections were stained with Alexa Fluor 488 anti-glutamine synthetase (#ab302584, Abcam, 1:100) and 647-conjugated donkey anti-mouse IgG (#715-606-150, Jackson ImmunoResearch, 1:100), along with the nuclear stain SYTO 83 (#S11364, Thermo Fisher, 1:10,000) for 1h at room temperature. Based on fluorescence signals, three phenotypically distinct regions were defined: GS + untreated, GS + CCl₄-treated, and IgG + CCl₄-treated. From each group, 20 regions of interest (ROIs) were selected (total, n=60) and subjected to UV illumination. The released RNA probes were collected for PCR amplification and purified using AMPure XP beads (#A63880; Beckman Coulter). Purified libraries were sequenced on the NextSeq 550 platform (Illumina) for spatial gene expression analysis.

### GeoMx data analysis

Raw FastQ data were converted to DCC files using GeoMx NGS Pipeline software (Version 3.1.1.6) and analyzed in R (version 4.5.1; R Core Team, R Core Team. 2024. R: Language and environment for statistical computing. Vienna (Austria): R Foundation for Statistical Computing. https://www.R-project.org/) using NanoString GeoMx Tools (version 3.21; Griswold M, Ortogero N, Yang Z, Vitancol R, Henderson D (2025). GeomxTools: NanoString GeoMx Tools. https://doi:10.18129/B9.bioc.GeomxTools, R package version 3.12.1, https://bioconductor.org/packages/GeomxTools) for quality control and normalization. Quality control assessment was performed for each ROI to evaluate the PCR contamination and sequencing depth. All 19,968 gene-associated probes passed the quality control. Of the original ROIs, the following passed quality criteria: GS+untreated (n=15), GS + CCl₄ (n=13), and IgG + CCl₄ (n=11). Following negative control normalization, gene expression data were analyzed in R (version 4.5.1) using the following packages: limma (version 3.64.3 according to (34)) for differential expression analysis, clusterProfiler (version 4.16.0 according to (35)) for gene set enrichment analysis, ggplot2 (version 4.0.0 according to (36)) for visualization, and ComplexHeatmap (version 2.24.1) for heatmap generation. For spatial deconvolution, we used SpatialDecon (version 1.18.0, according to (37)). which required an additional Q3 normalization of the ROIs.Reference cell type signatures were obtained from mouse single-cell RNA-seq data via the Celldex package (version 1.18.0; according to (38)), specifically using the MouseRNAseqData dataset based on (39, 40). Differential expression analysis between groups was conducted using the cutoff criteria of |log2FoldChange| > 1 and Benjamini-Hochberg adjusted p-value < 0.05, according to (41). Statistical significance was set at P < 0.05.

## Statistical analysis

All statistical analyses were performed using the GraphPad Prism v8. The relationship between quantitative IgG^+^ area (the novel marker) and established markers of liver injury (serum ALT, AST, and Ishak scores) was assessed using Spearman’s rank correlation coefficient (ρ). To assess the diagnostic accuracy of IgG^+^ area, ALT, AST, and Ishak scores, Receiver Operating Characteristic (ROC) curve analysis was performed. Significance between two groups was analyzed using Student’s t-test and between multiple groups using one-way ANOVA. Differences were considered statistically significant at p<0.05. Grubb’s test (extreme studentized deviate) was applied to determine whether extreme values were significant outliers. Data are represented as the mean ±SEM.

## Results

### IgG marks dead hepatocytes during acute liver intoxication and regeneration

To delineate the dynamics of hepatocellular injury and recovery, mice were exposed to a single high dose of carbon tetrachloride (CCl₄; 1600 mg/kg), and serum and liver tissues were collected in a time resolved (0, 1, 3, 6, 12, 24, 48, 72, 96, 144, and 384h; Fig. 1A) manner. Serum alanine aminotransferase (ALT) and aspartate aminotransferase (AST) levels increased from 1 h post-injection, reaching peak values at 24 h and 48 h, respectively, and peaked at 24 h (p < 0.001) and 48h (p < 0.01), respectively, reflecting extensive hepatocellular leakage (Fig. 1B and C). Thereafter, transaminase levels declined steadily, which was consistent with the onset of liver regeneration (Fig. 1B and C). Serial liver sections were stained with hematoxylin and eosin (HE) for histopathological evaluation (using Ishak necroinflammatory activity scores) and fluorescently labeled IgG to detect dead cells. Two independent pathologists assessed the HE-stained slides (Fig. 1D) and assigned Ishak scores. In line with ALT and AST kinetics, progressive necroinflammation, predominantly centrilobular, with maximal Ishak necroinflammatory activity scores was observed at 96h post-injury (Fig. 1E). ALT and AST levels are influenced by their respective half-lives and histological scoring remains semi-quantitative and dependent on expert interpretation, underscoring the need for robust surrogate markers. Therefore, IgG immunostaining was used as an alternative marker (Fig. 1D; Supp Fig. 1). Consistent with previous publications (27, 31, 42), IgG positivity was observed in sinusoidal vessels and in approximately 0.5% of hepatocytes, likely reflecting physiological turnover (Fig. 1F). Upon CCl₄ challenge, IgG positivity appeared at 6h, peaked between 72 and 96h, and nearly disappeared by 384h, precisely paralleling the biochemical and histological recovery phases (Fig. 1D). Quantitative analysis confirmed a strong correlation between the IgG-positive area and Ishak score (r=0.70, p<0.0001; Fig. 1G, left) and a moderate but significant correlation with serum AST (r=0.47, p=0.001; Fig. 1G, right). Receiver operating characteristic (ROC) analysis further demonstrated that IgG immunostaining exhibited the highest area under the curve (AUC=0.92) among all tested parameters (ALT, AST, Ishak score, AST/ALT ratio; Fig. 1H), establishing its superior diagnostic performance for quantifying hepatocellular death. To assess the molecular context of cell death, publicly available transcriptomic datasets from CCl_4_-treated mice (same strain and dosage; <u>E-MTAB-2445</u>) (33)) were analyzed. The expression of apoptosis- and necroptosis-related genes, including Casp3, Ripk3, Bak1, Casp2, Casp1, Bcl2, Bcl2l14, Bcl3, and Bcl10 was upregulated between 24-48h, corresponding to maximal IgG binding (Fig. 1I). This coordinated transcriptional activation substantiates that IgG binding encompasses both apoptotic and necrotic hepatocytes at the peak of injury, underscoring its mechanistic relevance and quantitative reliability as a biomarker of hepatocellular death.

**Fig. 1:**
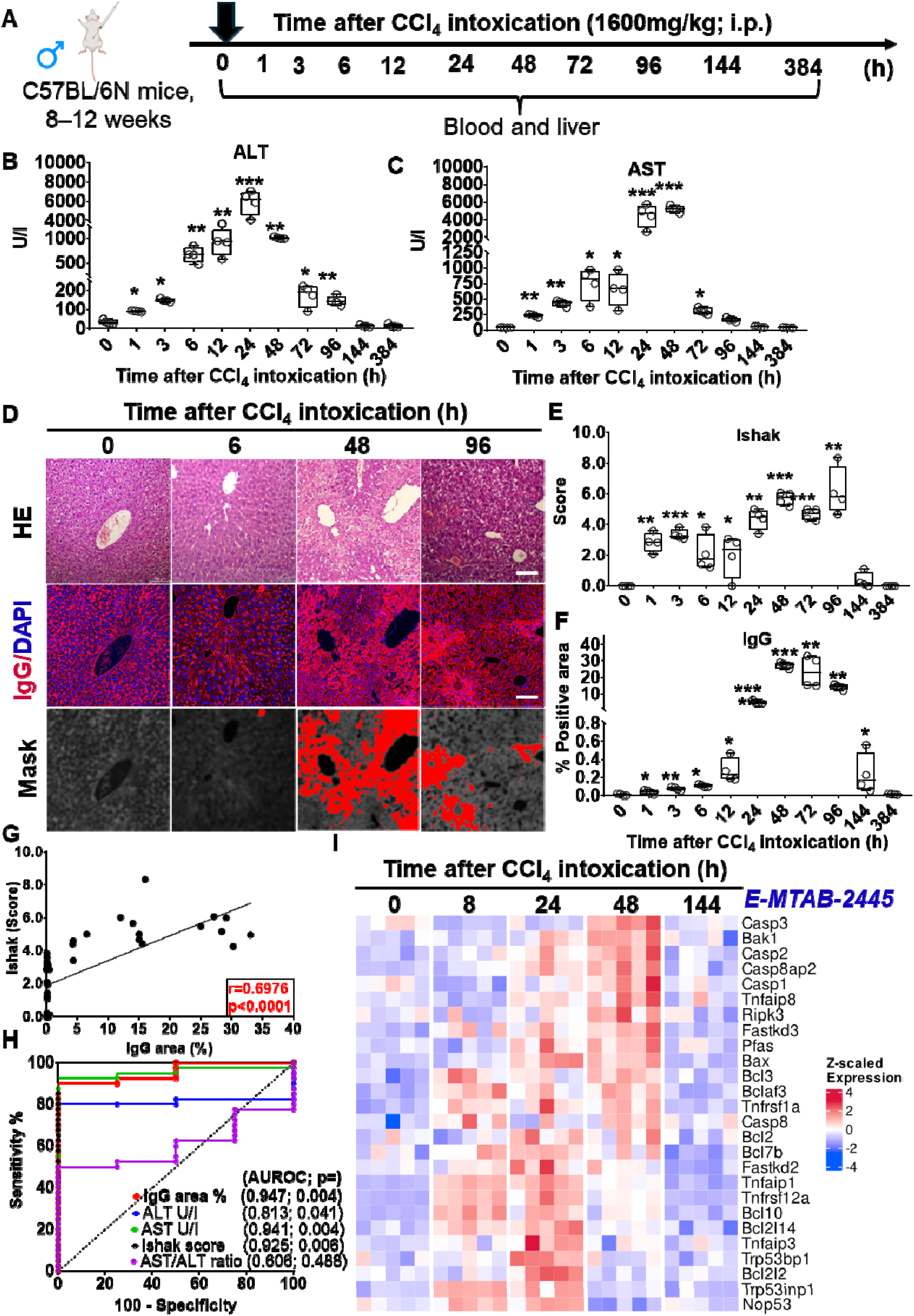
IgG staining identifies dead hepatocytes and is superior to ALT/AST for detecting CCl_4_-induced necrosis. (A) C57BL/6 mice received 1600 mg/kg CCl_4_; liver and serum collected across a 0-384 h time resolved manner. (B-C) Time course of serum ALT and AST as standard blood-based liver injury markers. (D) Representative serial sections: H&E (top), IgG (red) and DAPI (blue) immunofluorescence (middle), and IgG segmentation masks (bottom). Scale bars are 50 µm. (E, F) Quantification of necrosis by Ishak score (E) and IgG-positive area (F) over time. (G) IgG area strongly correlated with Ishak score (r=0.6976, p<0.0001). (H) ROC analysis: IgG area showed the highest AUC, demonstrating superior diagnostic accuracy for necrosis compared to ALT/AST and Ishak score. (I) Heatmap of programmed cell death genes (e.g., Casp3, Ripk3) upregulated at 24-48h (data from E-MTAB-2445 (33)), correlating with peak IgG staining. *p<0.05; **p<0.01; ***p<0.001.

**Supporting Fig. 1.**
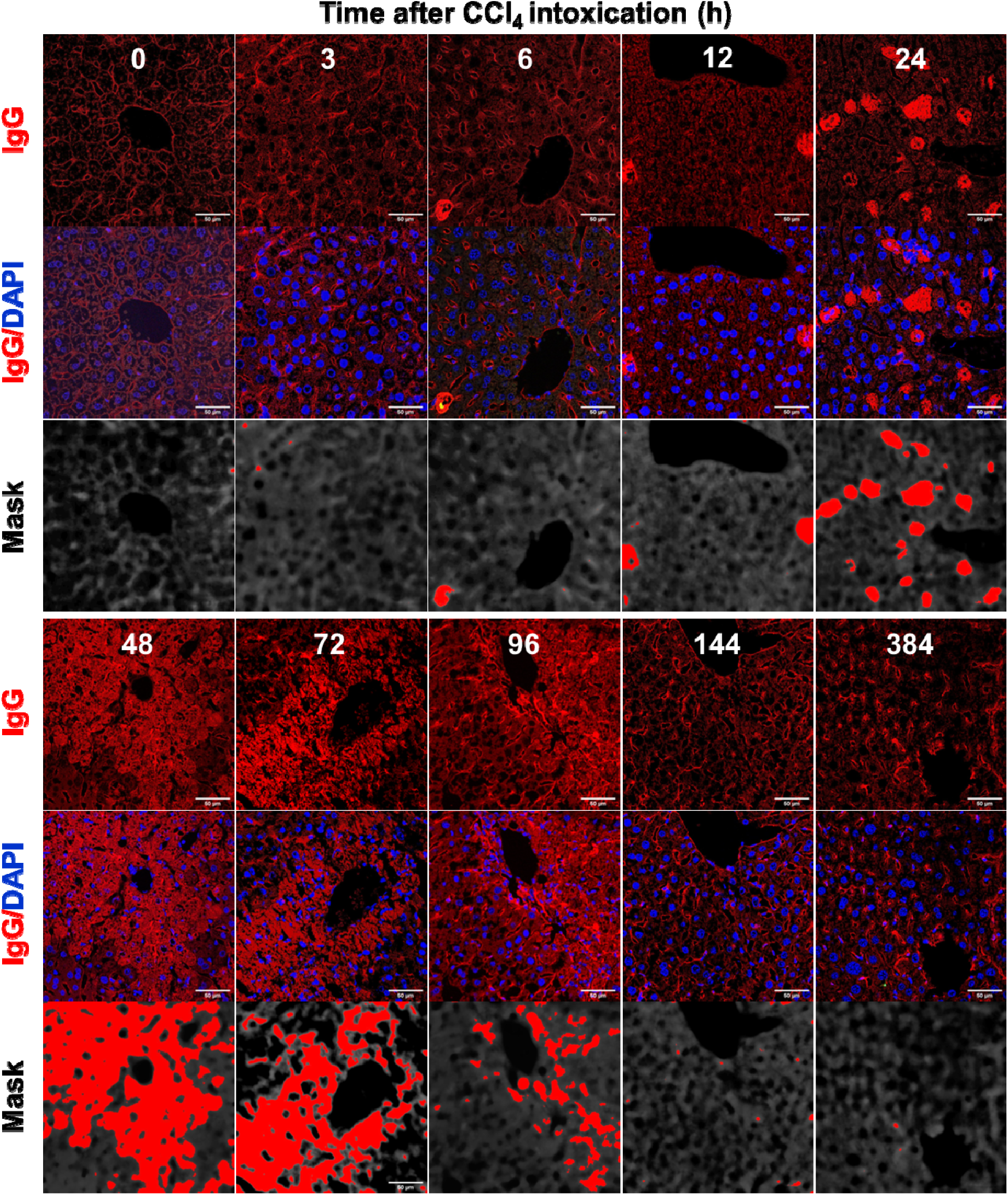
Time-resolved identification of dead hepatocytes by IgG staining during acute CCl₄-induced hepatotoxicity. Representative liver tissue sections at selected time points post-CCl₄ injection were stained with IgG (red) and DAPI (blue) (middle row) to visualize hepatocyte death. The corresponding IgG segmentation masks are presented in the bottom row. IgG-positive hepatocytes were first detectable 6h post-injection, reached a maximum at 48 h, and subsequently declined. Scale bars are 50µm.

### IgG positivity scales quantitatively with the severity of hepatocellular injury

To evaluate whether IgG labeling reflected the magnitude of hepatocellular injury, mice were administered increasing doses of CCl₄ (0, 25, 50, 100, 200, 400, and 800 mg/kg; n=4 per dose) and sacrificed 48h post-treatment, a time point corresponding to maximal histological damage (Fig. 2A). Serum ALT and AST levels exhibited a clear dose-dependent increase (Fig. 2B-C), confirming progressive hepatocellular leakage. Histopathology revealed corresponding centrilobular necrosis and inflammatory infiltration (Fig. 2D-E) as reflected by the Ishak score, which was assessed by two independent pathologists and reached its maximum at 400–800 mg/kg. IgG immunostaining of serial sections demonstrated dose-dependent increases in intensity and distribution, with low-dose exposure producing sparse pericentral labeling and high-dose exposure resulting in confluent necrotic zones spanning multiple lobules (Fig. 2D; Supplementary Fig. 2). Quantification revealed a strong positive correlation between the IgG-positive area and CCl₄ dose (Spearman ρ=0.719, p<0.0001; Fig. 2F). ROC curve analysis again identified the IgG-positive area as the most discriminative readout (AUC=0.988), outperforming the biochemical and histological metrics (Fig. 2G). Cross-parameter correlations demonstrated near-linear associations between IgG positivity and Ishak score (ρ=0.892), AST level (ρ=0.854), and ALT level (ρ=0.743; all p<0.0001; Fig. 2H). Collectively, these findings establish IgG labeling as a robust quantitative proxy for necrotic burden, accurately reflecting the intensity of hepatocellular damage in a pathologist-independent and reproducible manner.

**Fig. 2:**
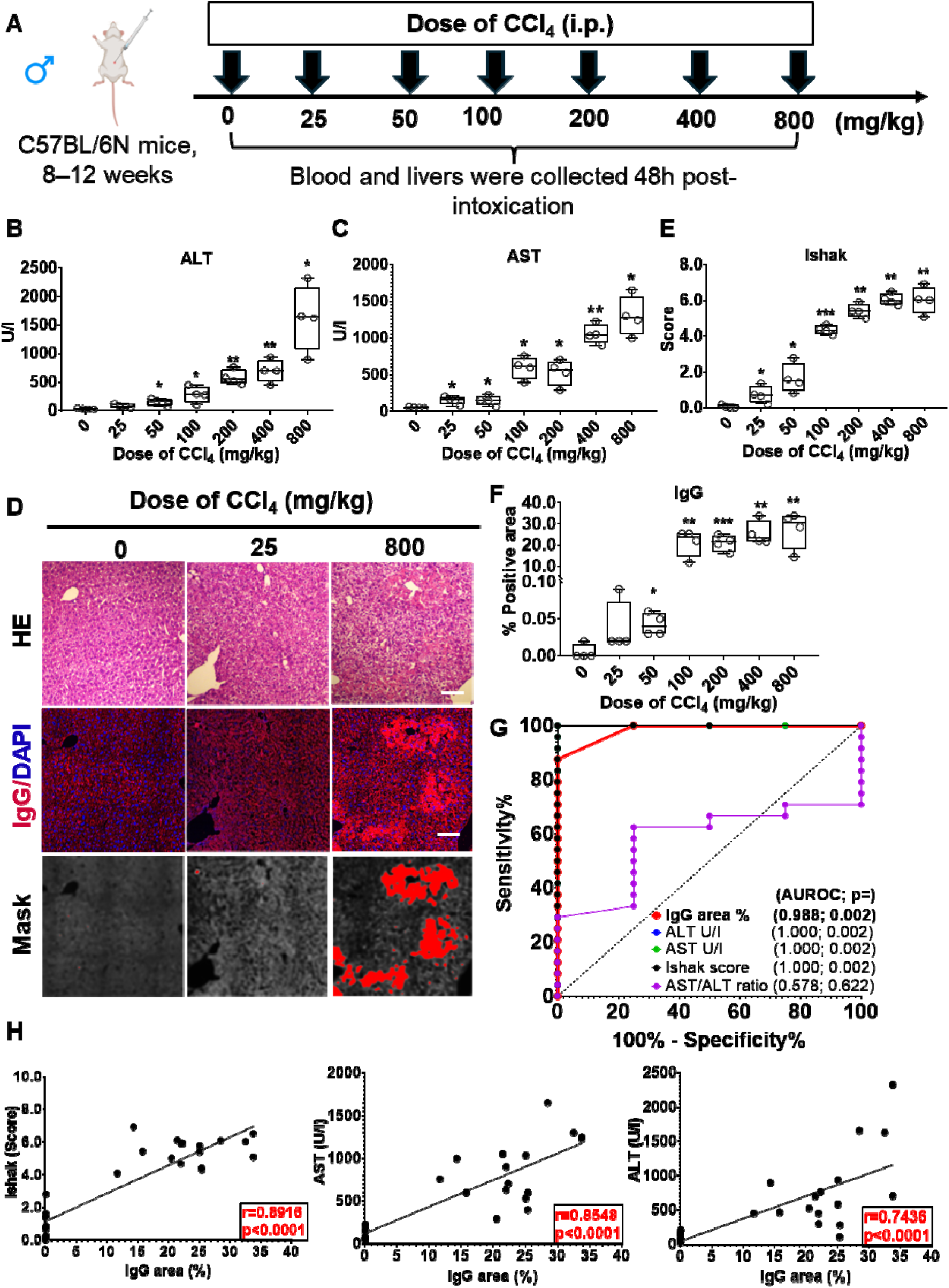
IgG area is a highly correlated and diagnostically superior marker of CCl_4_ dose-dependent liver necrosis. (A) Dose-response model: Mice (n=4/group) received increasing CCl_4_ doses (0-800 mg/kg) and were analyzed at 48h. (B-C) Serum ALT and AST levels demonstrated dose-dependent increases. (D) Representative H&E (top) and IgG (red) / DAPI (blue) immunofluorescence (middle) images show dose-dependent centrilobular necrosis, quantified via segmentation masks (bottom). Scale bars are 100µm. (E, F) Quantification confirms Ishak score (E) and IgG-positive area (F) increase robustly with CCl_4_ dose. (G) ROC analysis showed IgG area had the highest AUC, superior to all other markers. (H) Strong Spearman correlations were found between IgG area and Ishak score (r=0.892), AST (r=0.854), and ALT (r=0.743) (p<0.0001 for all).

**Supporting Fig. 2.**
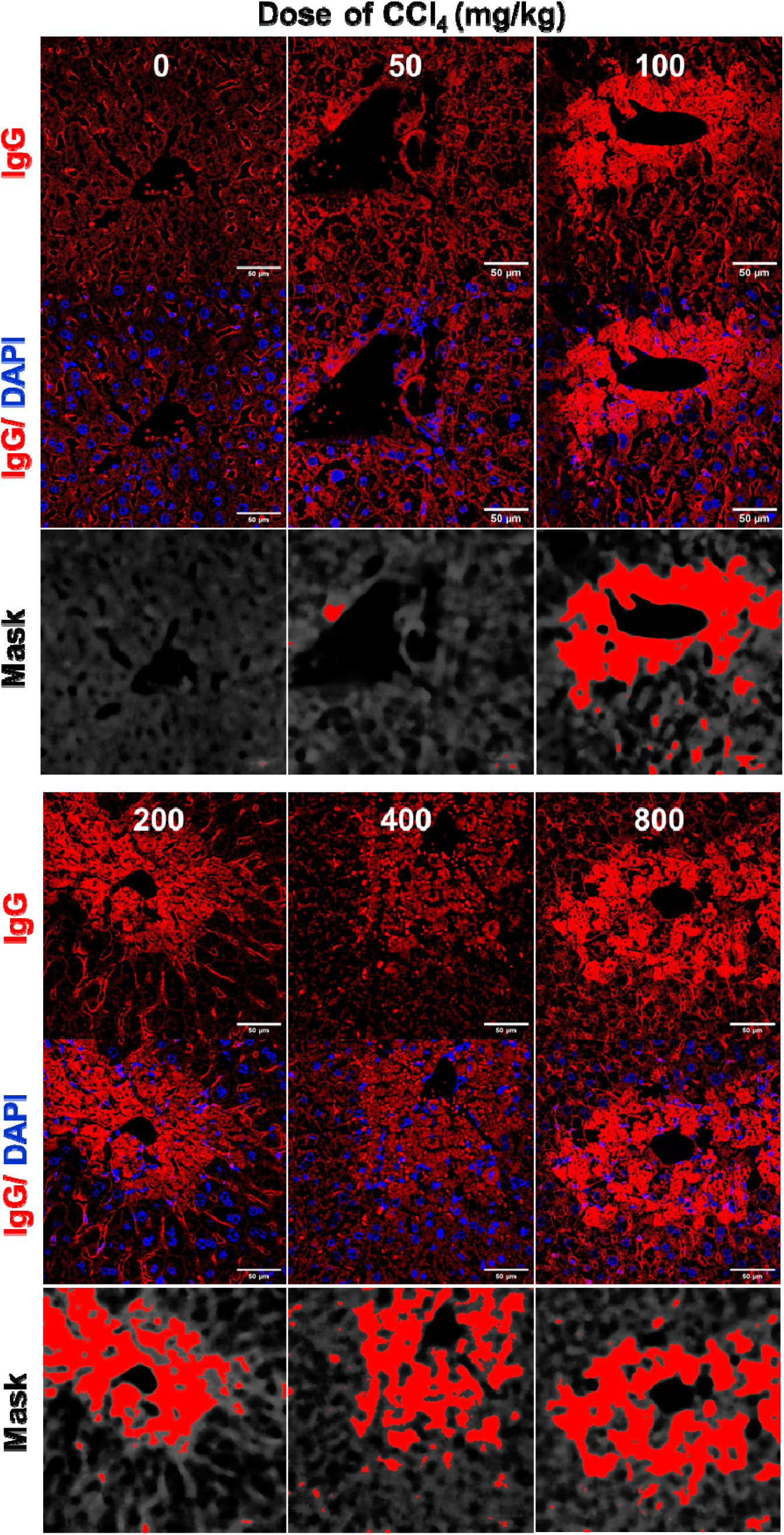
IgG staining reflects the extent of hepatocellular injury in a dose-dependent model of acute CCl₄-induced liver damage. Representative liver sections stained with IgG (red) and DAPI (blue) are shown in the middle row. IgG-positive areas expanded with increasing CCl₄ doses, particularly in centrilobular regions (400–800 mg/kg). The bottom row shows the threshold IgG signals used for the quantitative image analysis.

### IgG labels both necrotic and apoptotic hepatocytes

To delineate the mode of hepatocellular death recognized by IgG, dual immunofluorescence staining for IgG (red) and TUNEL (green) was performed at the time course following single CCl₄ administration. Control livers displayed negligible IgG or TUNEL staining, which is consistent with physiological turnover. In contrast, IgG-positive hepatocytes emerged as early as 24 h after post-intoxication, with a subset co-localized with TUNEL-positive nuclei (yellow/orange overlay; Fig. 3A-B; Supp Fig. 3), indicating the recognition of apoptotic cells with fragmented DNA. At 48-72 h, extensive co-staining was observed, corresponding to the peak phase of the hepatocellular injury. Quantitative image analysis revealed that IgG⁺TUNEL⁺ cells constituted approximately 20% of the total hepatocytes at 72 h, while IgG⁺TUNEL⁻ cells accounted for approximately 40%. The latter population likely represents necrotic remnants lacking nuclear DNA, reflecting IgG binding to exposed intracellular antigens within the disrupted membranes. These findings demonstrated that IgG recognizes epitopes generated during both apoptosis and necrosis, encompassing the full spectrum of terminal hepatocellular damage. Importantly, this dual recognition implies that IgG deposition integrates injury signals across distinct death pathways, providing a unified histological readout of cell loss, independent of morphological classification. IgG staining offers a powerful quantitative metric for assessing hepatocellular death kinetics, necrotic zone boundaries, and regeneration dynamics when applied to spatial and temporal injury mapping.

**Fig. 3.**
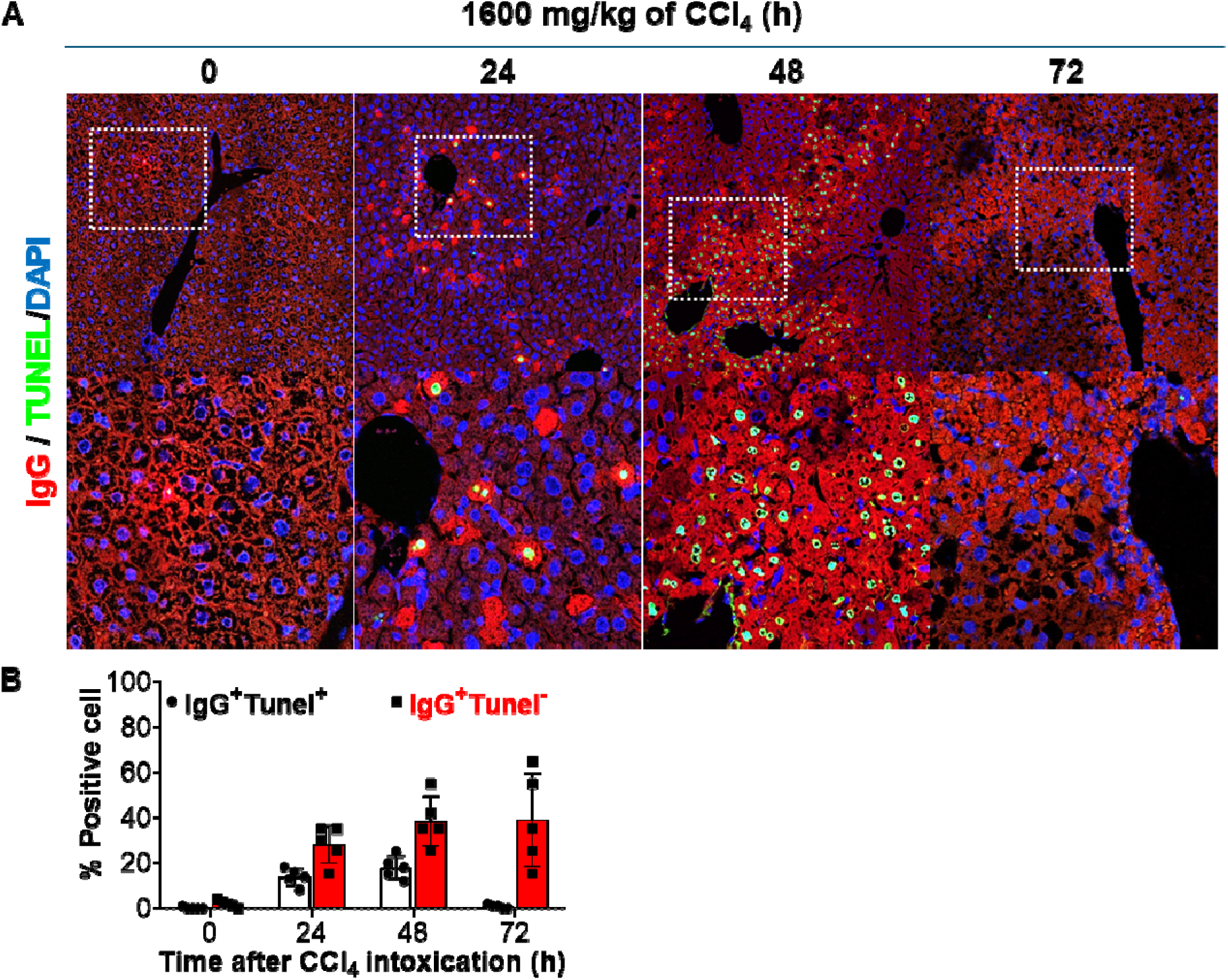
IgG labels both necrotic and apoptotic hepatocytes in the mouse liver following CCl₄ intoxication. (A) Immunofluorescence staining of liver sections from mice administered 1600 mg/kg CCl₄ at 0, 24, 48, and 72 h showed IgG (red) co-localizing with the apoptotic marker TUNEL (green) in some cells; nuclei were counterstained with DAPI (blue). (B) Quantification revealed that all apoptotic cells were IgG-positive (IgG⁺TUNEL⁺, black circles/bars), whereas others remained IgG-negative (IgG⁻TUNEL⁺, red squares/bars), and the proportion of IgG⁺ cells increased over time, reflecting progressive necrotic and apoptotic cell death after CCl₄-induced liver injury.

**Supporting Fig. 3.**
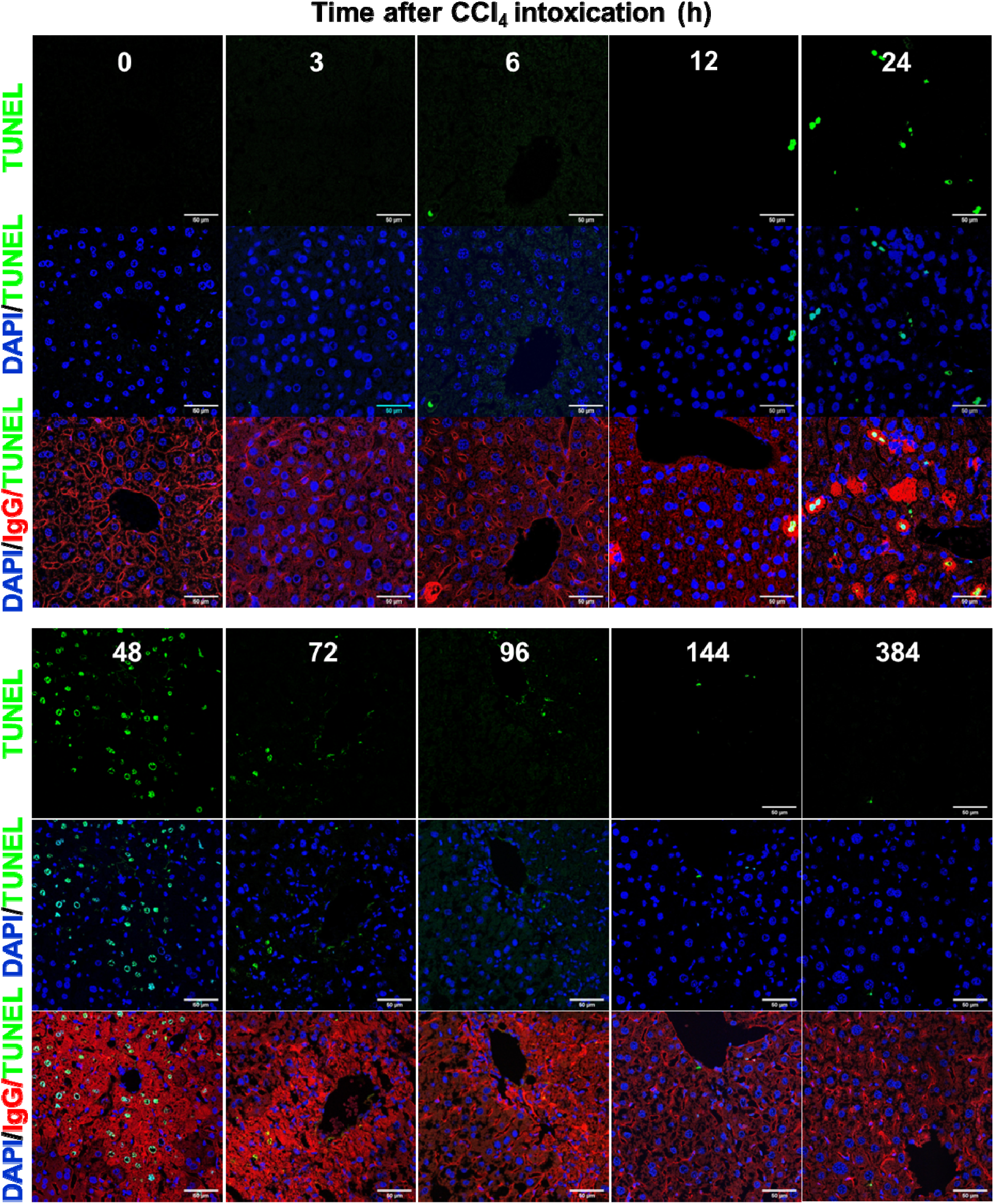
IgG labels both necrotic and apoptotic hepatocytes following CCl₄-induced liver injury. Liver sections from mice treated with 1600 mg/kg CCl₄ were stained for IgG (red), TUNEL (green), and DAPI (blue) over time. IgG colocalizes with some TUNEL-positive apoptotic cells. Quantification showed that all apoptotic cells were IgG⁺ (IgG⁺TUNEL⁺), whereas a subset remained IgG⁻ (IgG⁻TUNEL⁺). The proportion of IgG⁺ cells increased over time, reflecting progressive necrotic and apoptotic hepatocyte death after CCl₄ intoxication.

### IgG accumulates in dead hepatocytes independent of cell death etiology

We evaluated IgG deposition across multiple chronic murine liver injury models to assess the universality of IgG labelling in complex physiological contexts. These included a Western Diet (WD) model, WD combined with CCl₄-induced fibrosis, and cholestatic Mdr2−/− (hereafter Mdr2KO) injury, as well as the combined Mdr2-/-+CCl₄ model representing a mixed insult. IgG immunolabeling revealed distinct model-specific spatial patterns of deposition. In WD livers, IgG-positive hepatocytes (red signal) were scattered throughout the parenchyma, indicating diffuse metabolic stress (Fig. 4A). In contrast, WD+CCl₄ livers showed pronounced centrilobular IgG localization, reflecting combined metabolic and toxic injury centered around the central vein (Fig. 4A). This localized deposition was correlated with elevated serum transaminase levels (ALT and AST; Fig. 4B), indicative of ongoing hepatocellular injury. Quantitative assessment of in vivo IgG binding demonstrated a strong correlation with serum ALT and AST levels (Fig. 4B). ROC curve analysis confirmed that the IgG-positive area robustly discriminated between WD and WD+CCl₄ from SD groups, achieving the highest AUC compared to serum transaminases (Fig. 4C). In Mdr2-/- mice, IgG immunostaining predominated in the periportal and midzonal regions, consistent with cholestatic bile duct–associated injury (Fig. 4D; Supp Fig. 4). Finally, Mdr2-/-+CCl₄ livers displayed confluent panlobular IgG deposition, signifying extensive hepatocellular death spanning multiple lobular zones (Fig. 4D). This localized IgG deposition corresponded to elevated serum ALT and AST levels (Fig. 4E). Quantitative evaluation of IgG binding confirmed a strong correlation with transaminases, and ROC curve analysis demonstrated high diagnostic accuracy in distinguishing Mdr2-/- from Mdr2-/-+CCl₄ livers. Among all the tested parameters, the IgG-positive area consistently exhibited the highest AUC, underscoring its sensitivity and specificity as a marker of both chronic cholestatic injury and toxin-accelerated hepatocellular damage (Fig. 4F). These findings established that IgG accumulation reliably labels dead hepatocytes across diverse etiologies of liver injury, highlighting its utility as a universal in vivo marker of hepatocellular death.

**Fig. 4:**
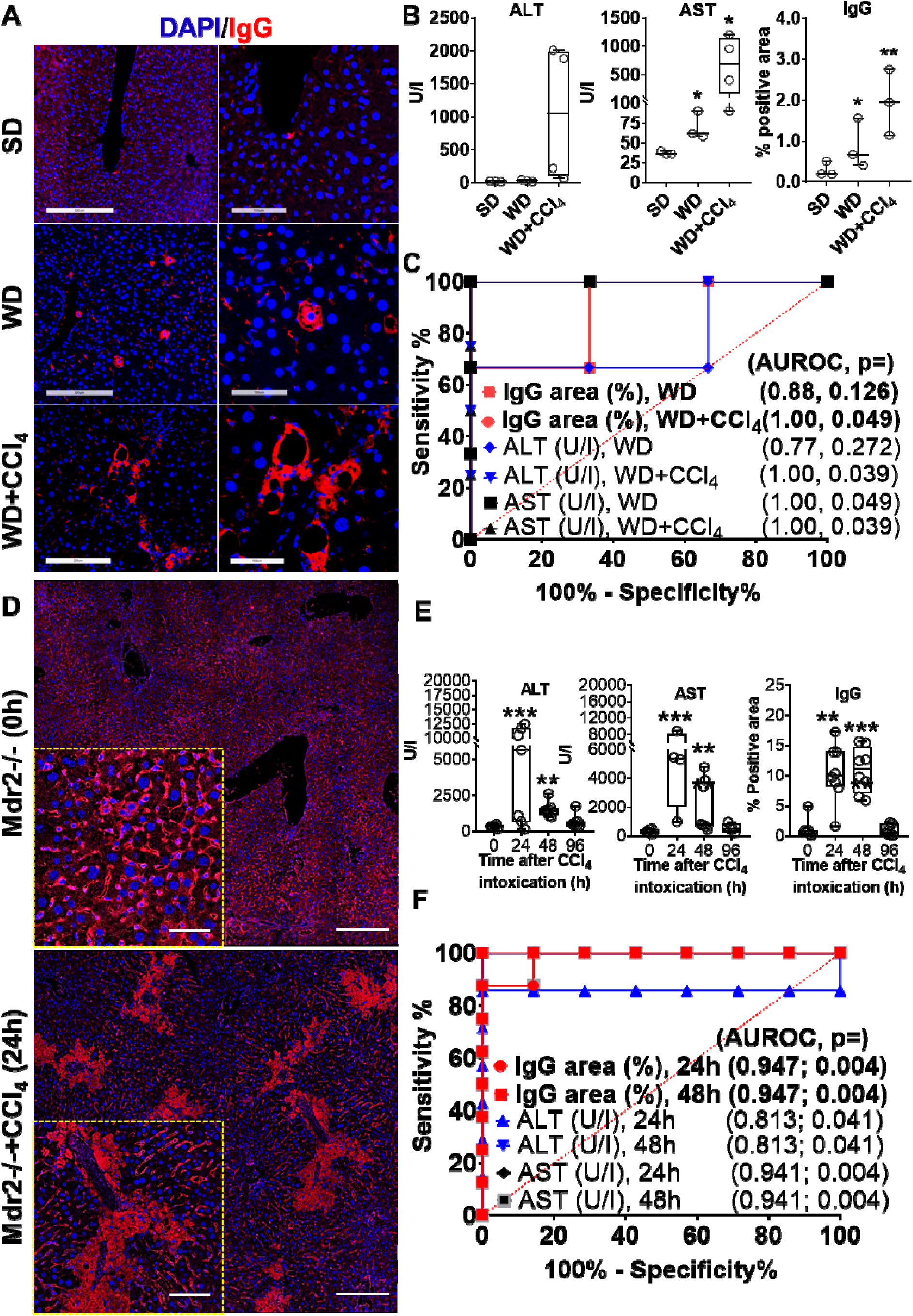
IgG deposition serves as a universal, pathologist-independent and spatial marker for hepatocellular death across diverse models of liver injury. (A) Immunofluorescence images of livers from standard diet (SD), Western diet (WD), and WD+CCl₄ mice stained for IgG (red) and DAPI (blue). WD livers showed scattered IgG⁺ hepatocytes, while WD+CCl₄ livers exhibited strong centrilobular IgG deposition. Scale bars are 300 and 100 µm in overviews and closeup images, respectively. (B) Serum ALT, and AST levels, as well as quantification of IgG⁺ area. (C) ROC analysis comparing IgG⁺ area with ALT/AST for distinguishing injury states WD or WD+CCl₄ vs. SD). (D) Immunofluorescence of Mdr2-/- and Mdr2-/-+CCl₄ livers. Mdr2-/- livers showed periportal/midzonal IgG staining, while Mdr2KO+CCl₄ showed panlobular IgG deposition. Scale bars are 300 and 50µm in overviews and close up images, respectively. (E) ALT, and AST levels, and IgG⁺ area, quantification confirmed strong positive correlations with disease progression. (F) ROC analysis showing IgG⁺ area outperforms or parallels ALT/AST for differentiating Mdr2-/- vs. Mdr2-/-+CCl₄ at 24 h and 48 h.

**Supporting Fig. 4.**
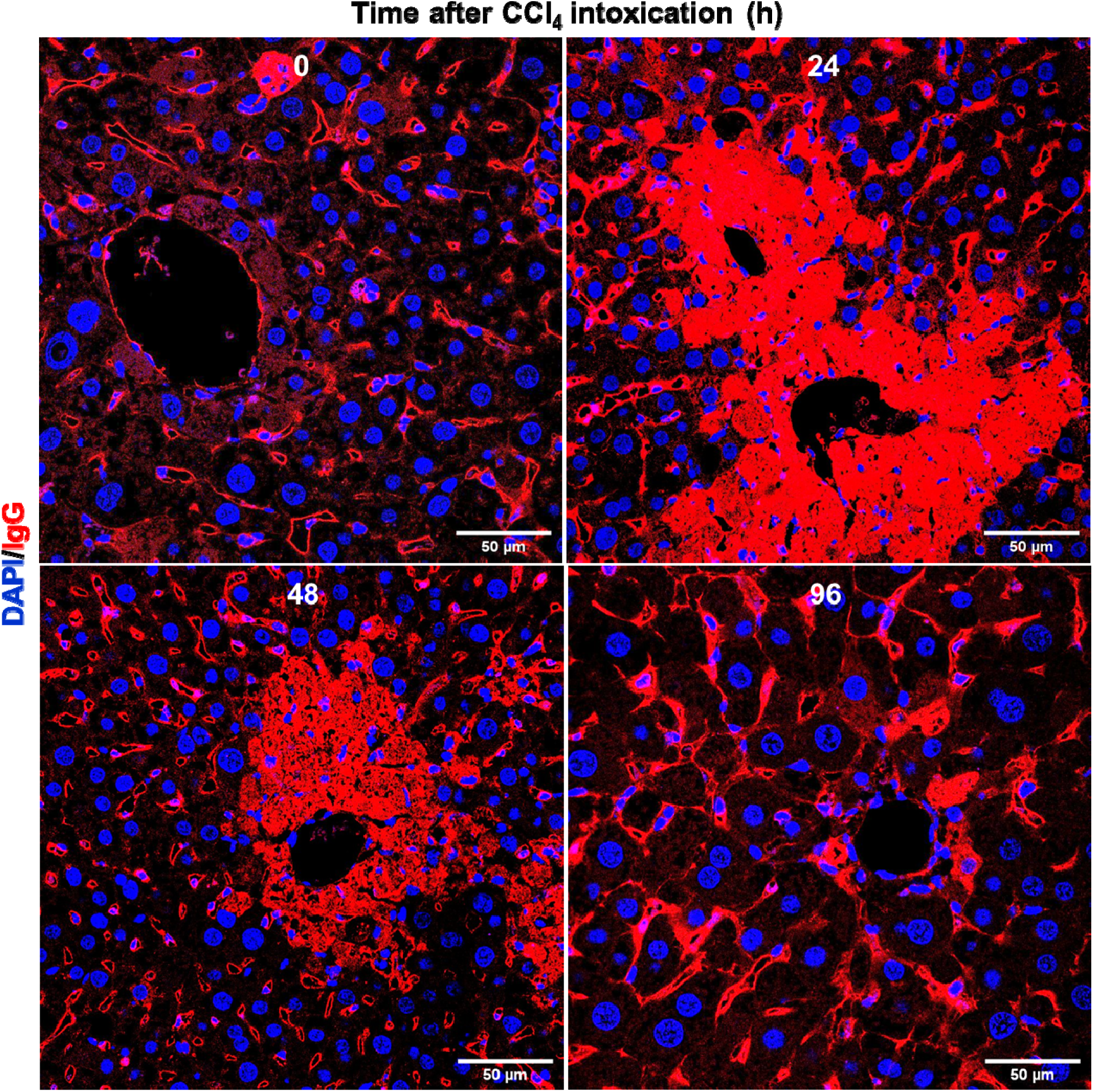
IgG deposition as a universal marker of hepatocellular death across acute and chronic liver injury models. Representative immunofluorescence images of Mdr2-/- (Mdr2KO) and Mdr2KO+CCl₄ liver sections stained for IgG (red) and counterstained with DAPI (blue). Mdr2KO livers exhibited periportal and midzonal IgG localization consistent with cholestatic injury, whereas Mdr2KO+CCl₄ livers displayed confluent panlobular IgG deposition, reflecting widespread hepatocellular death. Scale bars are 50µm.

### IgG positivity can serve as a spatial marker for necrotic lesions in combination with zonation, proliferation, and fibrogenic markers

To explore the spatial organization of necrosis, polarity, regeneration, and fibrogenesis, multiplex immunofluorescence staining was performed for IgG, polarity marker CD26, pericentral marker glutamine synthetase (GS), proliferation marker BrdU, and myofibroblast marker Acta2 (Fig. 5A–C). These analyses enabled the simultaneous visualization of hepatocellular death, zonal identity, proliferative activity, and fibrogenic responses within the same tissue sections. In untreated liver sections, IgG staining of hepatocytes was virtually absent, which was consistent with intact plasma membranes. In contrast, in injured liver tissue, IgG accumulation was observed as sharply demarcated zones of cytoplasmic fluorescence, predominantly in the hepatocytes adjacent to the central vein (Fig. 5A). The spatial confinement of IgG signals correlates with areas of hepatocyte necrosis and loss of membrane integrity, reflecting passive influx and retention of circulating immunoglobulins within damaged cells. Zonation analysis revealed that IgG deposition was localized to the GS-positive pericentral regions, indicating that necrosis occurred primarily within the central lobular domain (Fig. 5A), as expected. CD26-positive hepatocyte membranes were devoid of IgG signals and demarcated intact zones of viable tissues. The distinct spatial segregation between IgG and CD26 further confirmed the preservation of cell polarity, even under injury conditions (Fig. 5A), in the midzonal and periportal areas. Regenerative activity was assessed by BrdU incorporation, which identified proliferating hepatocytes predominantly in the peri-necrotic IgG-negative zones (Fig. 6B). These proliferative foci were localized adjacent to the GS-positive pericentral region, suggesting that hepatocyte regeneration originated from surviving cells at the necrosis boundary. Notably, no BrdU-positive nuclei were detected within the IgG-positive necrotic cores, indicating that IgG staining selectively marks terminally damaged or dead hepatocytes (Fig. 5B). Fibrogenic activation was visualized by α-smooth muscle actin (Acta2) immunostaining, which revealed the presence of Acta2-positive myofibroblasts within and encircling IgG-positive necrotic areas (Fig. 5C). This spatial proximity between dead hepatocytes and activated myofibroblasts indicates that IgG-defined lesions represent focal points of fibrotic initiation and wound-healing response. Collectively, these findings demonstrate that IgG immunostaining can be used as a robust marker for hepatocellular necrosis and membrane compromise. When integrated with additional markers such as GS for zonation, BrdU for proliferation, and Acta2 for fibrogenesis, IgG enables comprehensive spatial mapping of liver injury, regeneration, and fibrosis.

**Fig. 5.**
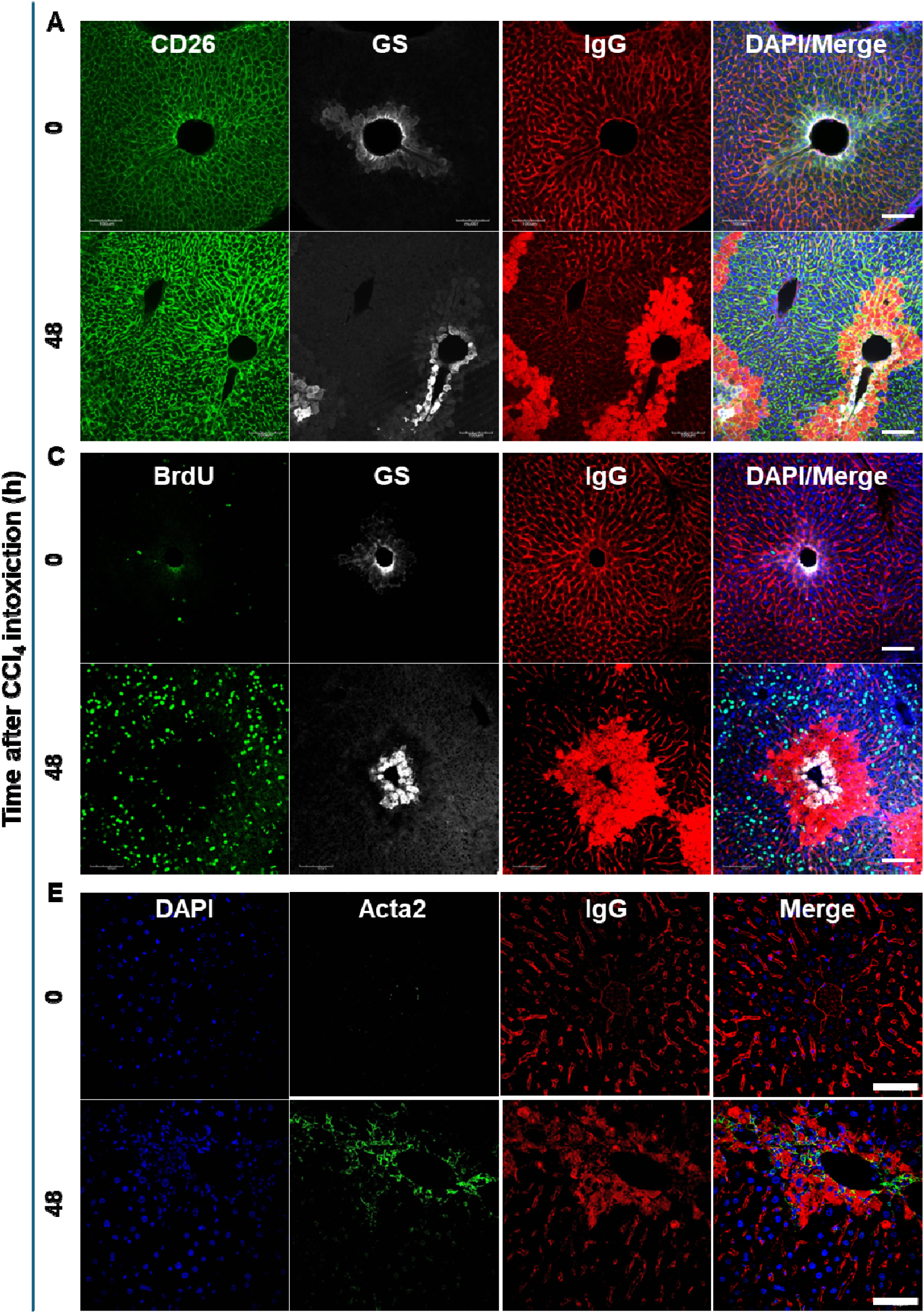
IgG accumulation defines necrotic hepatocytes and spatial injury zones within the hepatic lobule. (A) Co-immunostaining of IgG (red) with CD26 (green) and glutamine synthetase (GS, gray) delineates periportal and pericentral regions. IgG accumulation is confined to GS⁺ pericentral hepatocytes around the central vein, marking necrotic zones, while CD26 remains at the hepatocyte membrane, indicating preserved architecture. Scale bars are 100 µm. (B) Co-staining of IgG (red) with BrdU (green) and GS (gray) shows proliferating hepatocytes (BrdU⁺) localized to peri-necrotic, IgG⁻ regions, whereas IgG⁺ hepatocytes lack BrdU incorporation, confirming irreversible cell death. (C) Co-localization of IgG (red) with α-smooth muscle actin (Acta2, green) reveals activated myofibroblasts encircling IgG⁺ necrotic regions, indicating early fibrogenic remodeling. DAPI (blue) marks nuclei. Scale bars are 65µm.

**Fig. 6.**
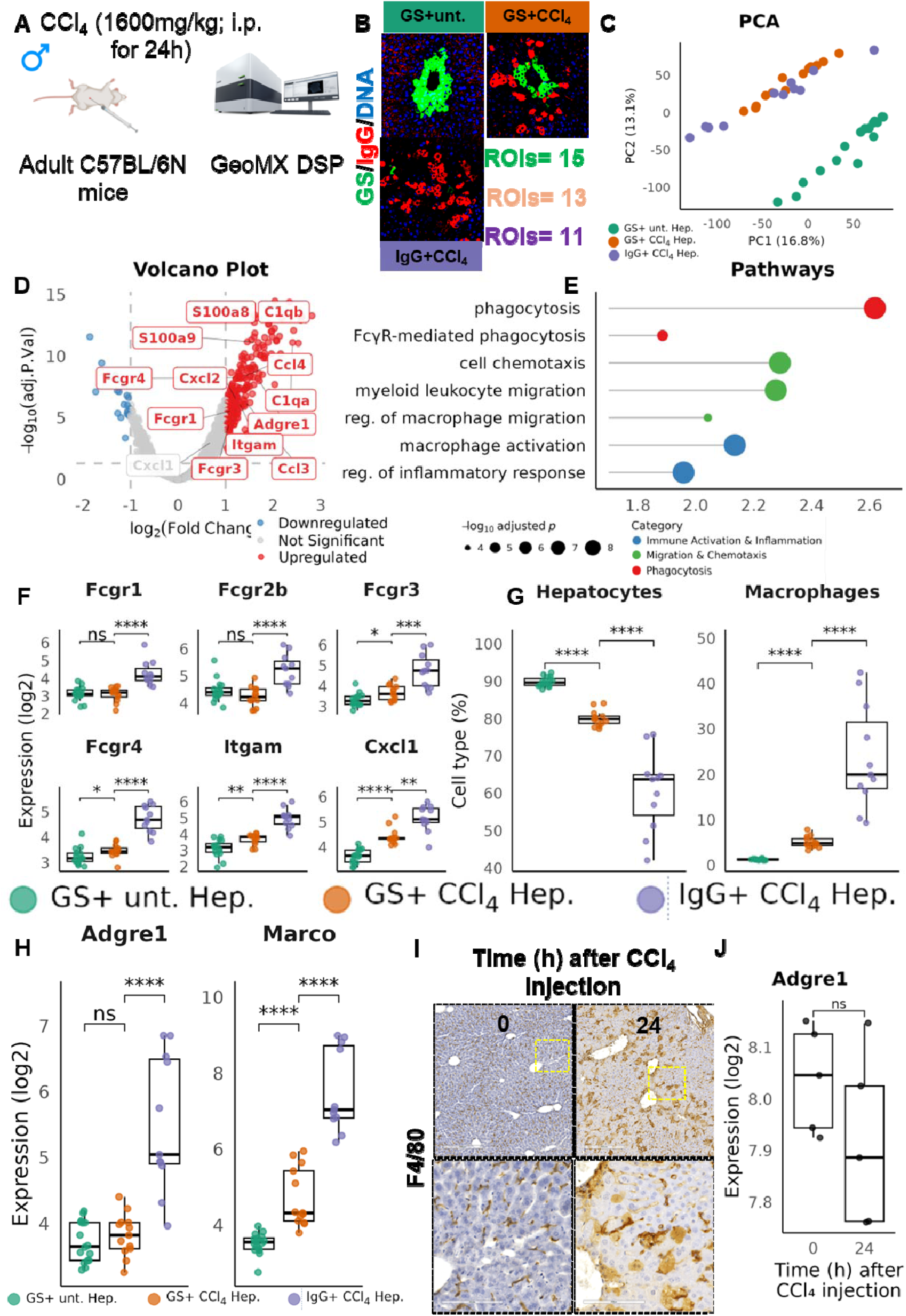
Molecular characterization of IgG⁺ hepatocytes in CCl₄-induced acute liver injury using spatial transcriptomics and immunohistochemistry. (A) Male C57BL/6N mice received a single CCl₄ injection (1600 mg/kg, i.p.; 24 h). Liver sections were analyzed using the GeoMx DSP Nanostring platform to spatially profile gene expression within defined Regions of Interest (ROIs). (B) GeoMx DSP images showing staining for GS (green), IgG (red), and DNA (blue). ROIs were selected from GS⁺ untreated hepatocytes (n=15), GS⁺CCl₄ hepatocytes (n=13), and IgG⁺CCl₄ hepatocytes (n=11). (C) PCA plot showing distinct clustering of the three ROI groups along PC1 (81% variance), indicating unique transcriptional profiles linked to injury and IgG deposition. (D) Volcano plot of differentially expressed genes between IgG⁺CCl₄ and GS⁺CCl₄ hepatocytes, highlighting significantly upregulated (red) and downregulated (blue) genes (adjusted p < 0.05). (E) KEGG and GO Biological Process enrichment analyses reveal increased immune activation, FcγR-mediated signaling, phagocytosis, chemotaxis, and macrophage migration/activation in IgG⁺CCl₄ regions. (F) Expression analysis shows significant upregulation of Fcgr1, Fcgr2b, Fcgr3, Fcgr4, Itgam, and Cxcl1 in IgG⁺CCl₄ ROIs. (G) Spatial deconvolution indicates IgG⁺CCl₄ ROIs contain higher macrophage fractions and fewer hepatocytes compared to uninjured controls. (H) *Adgre1* and *Marco*, well-established macrophage markers, were significantly upregulated in IgG⁺ CCl₄-treated ROIs. (I) IHC for macrophage marker F4/80 shows increased perivenous macrophage infiltration 24 h after CCl₄ intoxication (yellow dashed line). (J) Log₂-normalized Adgre1 (F4/80) expression remains unchanged at 24h post-CCl₄, indicating recruitment/migration without major transcriptional induction.

**Supplementary Fig. 5.**
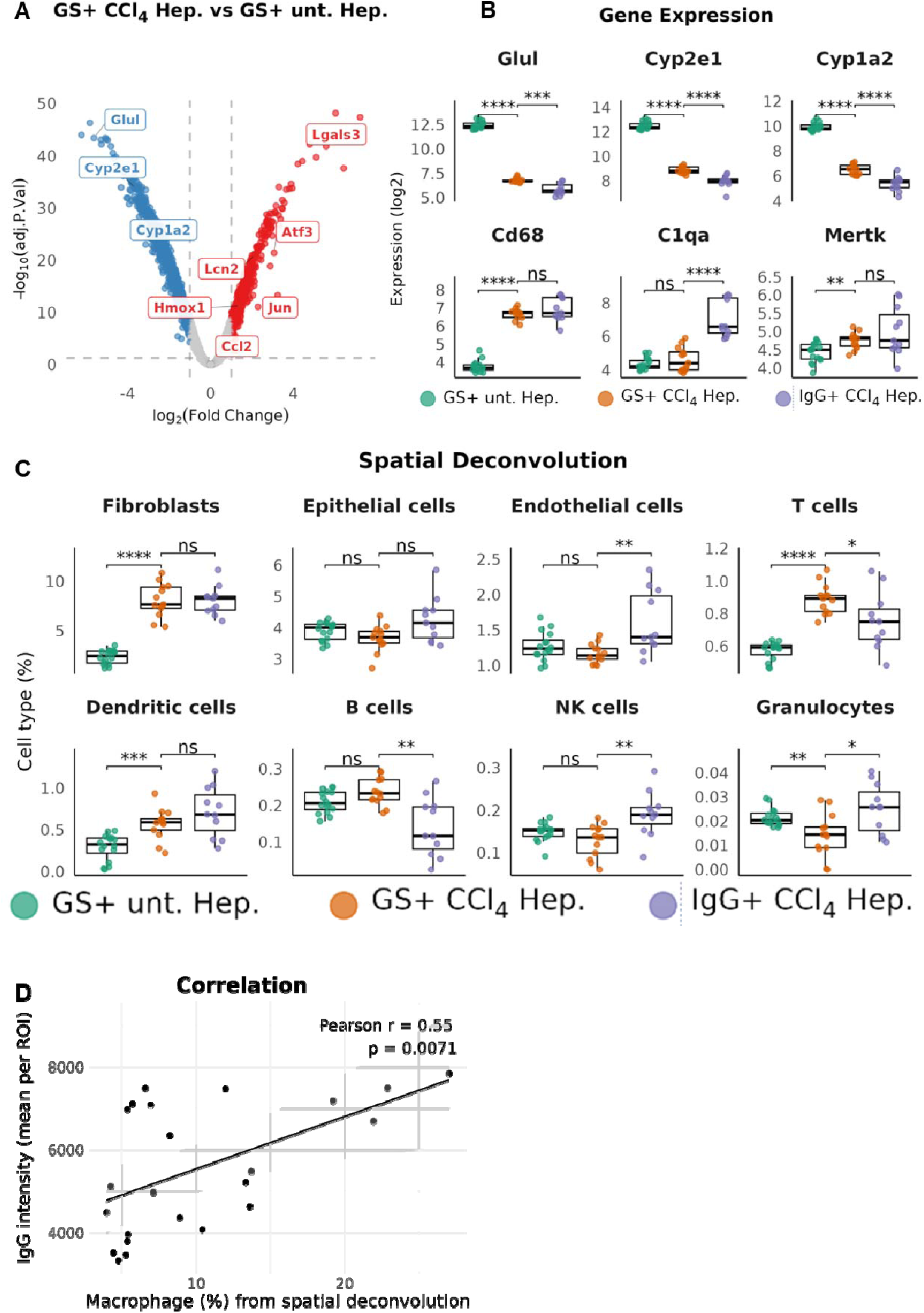
Spatial transcriptomic and cellular alterations in IgG⁺ hepatocytes following CCl₄-induced liver injury. (A) Volcano plot showing differentially expressed genes between GS⁺CCl₄ and GS⁺ untreated hepatocytes. Significantly upregulated genes are shown in red and downregulated genes in blue (adjusted p<0.05). (B) Log₂ expression of representative genes related to pericentral hepatocyte metabolism (Glul, Cyp2e1, Cyp1a2), macrophage markers (Adgre1, C1qa, Mertk), and inflammation across GS⁺ untreated, GS⁺CCl₄, and IgG⁺CCl₄ hepatocyte groups. (C) Spatial deconvolution analysis estimating proportions of immune and stromal cell types—including T cells, NK cells, granulocytes, fibroblasts, and others—across the same experimental groups. (D) Correlation plot showing a positive association between fibroblast abundance and a specified gene expression parameter (p = 0.007, R² = 0.55). Statistical significance: p < 0.05 (), < 0.01 (), < 0.001 (), < 0.0001 (****); ns = not significant.

### Spatial profiling reveals IgG-associated immune and macrophage cell infiltration and sinusoidal permeability in CCl_4_-treated livers

To investigate the molecular dynamics and characteristics of IgG-binding hepatocytes during the early phase of hepatotoxic injury, we performed spatial transcriptomic analysis on liver tissue collected one day after administration of CCl₄, a well-established model of pericentral (zone 3; glutamine synthetase (GS)) hepatocyte necrosis (Fig. 6A). Regions of interest (ROIs) were systematically selected based on multiplex immunofluorescence staining targeting immunoglobulin G (IgG), which marks necrotic or dying hepatocytes via antibody binding, and GS, which is a canonical marker of metabolically active pericentral hepatocytes. Based on spatial localization and marker expression, a total of 39 ROIs were classified into three distinct categories: (i) GS⁺ hepatocytes from untreated control livers, representing the physiological pericentral zone; (ii) GS⁺ hepatocytes from CCl_4_-treated livers, representing surviving hepatocytes within the injury-prone pericentral region; and (iii) IgG⁺ hepatocytes from CCl_4_-treated livers, corresponding to necrotic or IgG-tagged hepatocytes, indicative of immune complex formation and cell death (Fig. 6B). Principal component analysis (PCA) of the spatial transcriptomic data revealed distinct clustering of the two groups, with clear separation (with overlapping) between GS⁺ and IgG⁺ hepatocytes from CCl_4_-treated livers compared to GS^+^ untreated hepatocytes, highlighting transcriptional divergence in the context of injury (Fig. 6C). The volcano plot shows that IgG+CCl₄ Hepatocytes have significant upregulation of immune and inflammatory genes, including complement system, S100a8, Ccl3, Ccl4, Itgam (Cd11b), Fcgr1 and Cxcl1, compared to GS+CCl₄ ones (Fig. 6D; Supp Fig. 5A). This indicated that IgG positivity is associated with inflammatory and immune activation in CCl_4_-induced hepatic injury. To elucidate the molecular mechanisms driving these differences, we performed a pathway enrichment analysis. Compared to GS⁺ hepatocytes from untreated livers, both GS⁺ and IgG⁺ ROIs from CCl₄-treated samples showed marked upregulation of immune-related pathways, including phagocytosis, Fc gamma receptor (FcγR)-mediated phagocytosis, cell chemotaxis, myeloid leukocyte migration, regulation of macrophage migration and activation, as well as inflammatory response pathways (Fig. 6E), reflecting robust inflammatory activation and local vascular remodeling in the injured tissue, as suggested by Mattos et al. (27). Next, we examined the expression of Fcγ receptors and chemokines (Fig. 6F). We reported a significant upregulation of Fcgr1, Fcgr2b, Fcgr3, Fcgr4, Itgam, and Cxcl1 in IgG+CCl₄ ROIs, consistent with the findings of Mattos et al. (27). As expected, pericentral functions, that is, Glul, Cyp2E1, and Cyp1a2, were downregulated; however, macrophage markers were upregulated (Suppl. Fig. 5B). Using computational deconvolution of the cell-type signatures, we quantified the relative abundance of hepatic cell populations within each ROI. T cells, NK cells, granulocytes, and fibroblasts (Supp. Fig. 5C) were significantly enriched in ROIs from CCl_4_-treated livers, particularly within IgG⁺ regions, whereas hepatocyte fractions were notably decreased, and macrophages increased, reflecting immune cell infiltration into necrotic zones (Fig. 6G). A particularly striking finding emerged when IgG transcript levels were examined across the ROIs. There was a significant positive correlation between IgG expression and the number of macrophage-rich ROIs per sample (R²=0.65, p=0.007; (Suppl. Fig. 5D), suggesting that macrophage accumulation was closely associated with local IgG abundance. This may reflect the FcγR-mediated engagement of macrophages with IgG-opsonized cellular debris or immune complexes. This is confirmed by upregulation of well know macrophage markers namely Adgre1 (encoding the macrophage marker F4/80) and Marco (a marker for sessile macrophages) in IgG+CCl4 ROIs (Fig. 6H). Immunohistochemical staining for F4/80 in the intoxicated liver showed macrophage infiltration into necrotic regions (Fig. 6I). To further validate this observation, transcriptomic analysis of publicly available datasets (E-MTAB-2445; (33)) revealed no significant upregulation of Adgre1 one day after CCl₄ treatment, implying that the macrophages identified in IgG⁺ zones likely migrated into the tissue rather than being locally expanded (Fig. 6J). Collectively, these spatially resolved datasets support a model in which IgG-positive hepatocyte zones act as immunological hotspots in acutely injured livers. These areas facilitate the recruitment and retention of IgG-reactive immune cells, particularly macrophages and granulocytes, likely via a combination of FcγR engagement and chemokine-mediated trafficking. These findings suggest that either IgG–FcγR interactions or modulation of local vascular permeability may explain IgG binding to dead cells as shown in graphical abstract.

## Discussion

Our findings established IgG immunostaining as a reliable, sensitive, pathologist-independent, and reproducible marker of hepatocellular injury in both acute and chronic liver disease models. Mechanistically, the accumulation of IgG within necrotic hepatocytes reflects its binding to ubiquitous intracellular and membrane-associated components, including DNA, fibrin, actin, tubulin, lysophosphatidylcholine, and oxidized phospholipid debris (43–45). Traditionally, ruptured and intact plasma membranes delineate necrotic and apoptotic cell death, respectively, allowing serum immunoglobulins to infiltrate the cytoplasm during necrosis (46). Our immunostaining approach directly capitalizes on this physiological influx of endogenous IgG, effectively utilizing the host’s own circulating antibodies as intrinsic markers of membrane loss. This process represents the morphological endpoint of hepatocellular death and provides a spatially resolved correlation with the biochemical indicators of injury. Notably, the temporal emergence of IgG-positive hepatocytes closely paralleled classical biochemical and histological parameters, namely serum ALT and AST activities and Ishak necroinflammatory scores (15), thereby confirming strong concordance among biochemical, histopathological, and immunohistochemical indicators (7, 32, 47).

While conventional injury metrics, such as ALT/AST quantification and semi-quantitative histological scoring, remain widely used, they have inherent limitations, including indirect readouts, delayed kinetics, and observer-dependent variability (10, 48–51). Moreover, aminotransferases lack absolute liver specificity, as extrahepatic sources such as skeletal muscle can confound interpretation (52, 53). In contrast, IgG immunolabeling provides a direct morphological signature of hepatocyte death. This approach minimizes subjectivity and enhances reproducibility when combined with automated image quantification. In dose-dependent CCl₄ models, IgG deposition scaled with injury severity and was localized precisely to centrilobular necrotic zones, correlating with biochemical and histological indices of tissue damage (31, 32). These characteristics underscore the potential of IgG staining as a quantitative and high-throughput biomarker for toxicological screening and predictive modeling of hepatocellular death and regeneration dynamics.

Interestingly, IgG labeling was detected in both apoptotic and necrotic hepatocytes, colocalized with TUNEL positivity, thereby capturing both passive and active modes of cell death. Natural antibodies recognize intracellular antigens, such as DNA, histones, and actin, which are exposed after membrane disruption, facilitating opsonization and subsequent clearance by phagocytes (27, 54). Consequently, IgG deposition functions as an “eat-me or find-me” signal, linking humoral recognition to innate effector mechanisms. Recent studies have shown that complement components (C1q and C3) co-deposit with IgG on necrotic hepatocytes, enhancing macrophage and neutrophil phagocytosis, whereas disruption of this process delays debris clearance and impairs regeneration (55, 56). Thus, IgG immunostaining not only demarcates necrotic hepatocytes but also reflects the initiation of an immunologically coordinated tissue response for repair. Complementary pulse–chase assays using Trypan Blue corroborate that IgG-positive cells correspond to membrane-compromised hepatocytes, validating the specificity of this marker in vivo (42).

Multiplex immunofluorescence further revealed that IgG-positive necrotic regions were consistently bordered by BrdU^+^ proliferating hepatocytes and infiltrated by Acta2^+^ myofibroblasts (7, 32, 57, 58). This perinecrotic organization illustrates how hepatocellular death generates localized instructive cues. These cues stimulate hepatocyte proliferation to restore parenchymal integrity while concurrently activating fibroblasts and stellate cells to deposit the extracellular matrix (59), thereby maintaining wound edges. These findings emphasize the modularity of liver regeneration, suggesting that spatially confined injury orchestrates parallel regenerative and fibrogenic programs through locally derived rather than systemic signaling, as documented by others (60–62). Quantitative mapping of necrotic and proliferative compartments thus provides valuable input for computational models of regeneration-fibrosis balance and for evaluating pharmacological interventions targeting this dynamic equilibrium (7).

Spatial transcriptomic analyses further supported the immunological dimension of IgG-marked injury zones. These regions are enriched in macrophages and granulocytes and display heightened Fcγ receptor signaling, phagocytosis, and chemokine (Ccl2 and Cxcl1) pathway activation (27). Such molecular signatures indicate that macrophage accumulation occurs primarily through recruitment rather than proliferation, establishing IgG deposition as both a morphological and immunological hotspot of liver injury. The interplay between FcγR engagement and cytokine release likely shapes the transition from inflammation to repair, a mechanism that may underlie divergent outcomes in acute and chronic injury contexts.

A limitation of the present study was the focus on preclinical models, which may not fully capture the heterogeneity of human liver pathology. Future studies should evaluate IgG staining in well-characterized patient cohorts and integrate it with digital pathology pipelines for clinical validation. Additionally, mechanistic exploration of Fcγ receptors and complement signaling in human samples may identify therapeutic strategies to modulate inflammation and enhance tissue repair, as suggested by Mattos et al. (27).

## Conclusion

In summary, this study posits IgG immunostaining as a mechanistically informed, spatially precise, pathologist-independent, and translationally versatile marker of hepatocellular death. By integrating biochemical, histological, and immune perspectives, IgG labeling enables high-resolution mapping of liver injury, regeneration, and fibrogenesis. The convergence of immunohistochemistry with multi-omics and computational tissue modeling offers a promising framework for monitoring therapeutic efficacy, predicting regenerative capacity, and identifying immunologically active microenvironments in liver diseases.

## ACKNOWLEDGEMENT

The authors acknowledge support from the Core Facility for Imaging and Microarray Unit Mannheim, Medical Faculty Mannheim, Heidelberg University.

## FUNDING

This study was supported by the German Federal Ministry of Education and Research (BMFRT) through the LiSyM Project (Grant Numbers: PTJ-FKZ 031L0043, 031L0257A, and 031L0314A), LiSyM cancer Bode: FKZ: 031L0257G, RHEACELL GmbH (Grant Number: 72000069), and Stiftung für Biomedizinische Alkoholforschung (Grant Number: 73000350).

## AUTHOR CONTRIBUTION

LN, PE, AD, AO, JW, JA, BB, IvR, TC, SW, and SH conceived the animal studies, blood analysis, IHC and IF staining, and image analysis. LN, AD, and SH performed spatial transcriptomics. LN, AD and SH wrote the manuscript. ME, JB, JH, and SD performed critical revision of the manuscript. SH and SD acquired funding for this study. All authors have read the final version of the manuscript.

## Notes

### Competing Interest Statement

The authors have declared no competing interest.

